# Application of self-organizing maps to AFM-based viscoelastic characterization of breast cancer cell mechanics

**DOI:** 10.1101/2022.12.03.518961

**Authors:** Andreas Weber, Maria dM. Vivanco, José L. Toca-Herrera

**Author notes:** **Contact Information:** (current address).

## Abstract

Cell mechanical properties have been proposed as label free markers for diagnostic purposes in diseases such as cancer. Cancer cells show altered mechanical phenotypes compared to their healthy counterparts. Atomic Force Microscopy (AFM) is a widely utilized tool to study cell mechanics. These measurements often need skilful users, physical modelling of mechanical properties and expertise in data interpretation. Together with the need to perform many measurements for statistical significance and to probe wide enough areas in tissue structures, the application of machine learning and artificial neural network techniques to automatically classify AFM datasets has received interest recently. We propose the use of self-organizing maps (SOMs) as unsupervised artificial neural network applied to mechanical measurements performed via AFM on epithelial breast cancer cells treated with different substances that affect estrogen receptor signalling. We show changes in mechanical properties due to treatments, as estrogen softened the cells, while resveratrol led to an increase in cell stiffness and viscosity. These data were then used as input for SOMs. Our approach was able to distinguish between estrogen treated, control and resveratrol treated cells in an unsupervised manner. In addition, the maps enabled investigation of the relationship of the input variables.

## Introduction

Mechanical forces are important for cell migration, interaction of cells with their environment, tissue morphogenesis and in disease ^1–6^. In cancer, due to deposition of aligned extracellular matrix proteins solid, stiff tumours can be recognized on the macroscale by palpation and similar techniques ^7^. Interestingly, on the single cell scale, cancer cells have been found to exhibit a softer, more fluid like phenotype ^8^. This change is conserved over many types of cancer tissue origin, ranging from breast, brain, prostate, kidney to lung cancer. Besides other biochemical alterations in cancer cell metabolism, they are thought to have a more disorganized cytoskeletal network which contributes to their mechanical phenotype ^9^. In addition, the softening of cells appears to correlate with the level of aggressiveness, as well as changes in the adhesive and motile properties of these cells ^10–12^. Therefore, the mechanical phenotype of cancer cells is thought to be a promising, label-free biomarker ^13–16^. Different studies have provided evidence for the softening of cancer cells and tissue, as well as enabled the differentiation of cancer progression in tissue sections.

Mechanical properties of cells and tissue can be measured either by active or passive methods ^17,18^. The first include methods such as parallel plate rheometry, optical and magnetic tweezers, micropipette aspiration, optical stretcher, magnetic twisting cytometry and AFM. AFM is a widely applied technique, enabling imaging with nanometric precision while measuring mechanical properties at a wide set of forces, strains, and frequencies. In AFM measurements, the probe (e.g., a micrometric particle or a nanometric tip) is brought into contact with the sample and the bending of the cantilever is measured. The bending in relation to the position and stiffness of the cantilever can be converted to so called force-distance and force-time curves. By application of different contact mechanics (elasticity, viscoelasticity), mechanical properties of the sample are determined ^19^. Sample handling, performance of the measurements and data fitting routines often need expert knowledge. There is the notion that AFM based single cell and tissue mechanics could be applied in clinical settings to support medical professionals in classification of samples such as cancer progression through evaluation of biopsies ^20,21^.

Recently, machine learning and neural network algorithms have been applied to surface probe microscopy data ^22^, including image segmentation ^23^, automatic data processing ^24–27^, cancer cell classification and progression grade evaluation ^28–32^. All these approaches were performed in a supervised way, consisting of a training set with defined classification, that could be used to classify data after training. In this work, we apply self-organizing maps (SOMs, also called Kohonen maps) as an artificial neural network approach to analyse mechanical measurements of breast cancer cells ^33–36^. SOMs are unsupervised artificial neural networks that enable 2D visualization of multidimensional data space while still conserving the topology of the dataset. They have been widely applied in data science, speech processing, and recently also in life sciences and chemometrics. One result of SOMs are U-matrix plots that show the similarity/dissimilarity between neurons, enabling post hoc cluster analysis.

Breast cancer is the most commonly diagnosed cancer and the first cause of death from cancer in women worldwide ^37^. A common therapeutic approach in hormone-dependent breast cancer, the most common type of breast cancer, is to target estrogen receptor (ER) signalling with drugs such as tamoxifen or aromatase inhibitors. ER signalling is widely recognized in playing active roles in cytoskeleton, motility and adhesion protein expression with multiple downstream targets ^38,39^. The alterations in the mechanical phenotypes of breast cancer cells treated with drugs that either inhibit or activate ER can be studied *in vitro* ^40,41^.

In this work we have used epithelial breast cancer (MCF-7) cells as model for breast cancer cell mechanics and performed stress relaxation measurements via AFM using sharp pyramidal tips. We then applied a viscoelastic model to derive mechanical parameters that were used as input layer for a SOM, which was followed by cluster analysis. We show the principal ability of SOMs to differentiate between control, estrogen and resveratrol treated cells, while the SOMs placed control and tamoxifen treated cells in close neighbourhood.

## Results

### Estrogen receptor interacting drugs change breast cancer viscoelastic properties

Table 1 shows the values derived from the 5-element Maxwell model fitting for the stress relaxation segments of the cells measured via AFM after the three different treatments (plus the carrier, as control). Control cells showed moduli in the range of a few hundred Pa and relaxation times of 0.2 and 3.1 s respectively. Treatment with estrogen led to significant softening and less viscous mechanical phenotype, reproducing other published results ^41,42^. The treatment with tamoxifen did not lead to significant changes in the mechanical properties of the cells, according to this analysis. Finally, treatment with resveratrol led to a stark increase of moduli of up to 6 times, slightly longer relaxation times and threefold increased viscosities. These data overall agree well with published literature ^40^. Note that some of the parameters showed strong correlations, which will be discussed at a later point.

**Table 1.** Viscoelastic properties of the different groups of treated cells derived from the stress relaxation measurements. The values show mean value ± standard error of mean.

| | Control | E2 (100 nM) | TAM (10 μM) | RESV (50 μM) |
| --- | --- | --- | --- | --- |
| $\delta$ [μm] | 2.11 ± 0.05 | 2.82 ± 0.05 | 2.04 ± 0.04 | 1.32 ± 0.05 |
| $E_{\infty}$ [Pa] | 444 ± 23 | 187 ± 11 | 510 ± 25 | 2959 ± 235 |
| $E_1$ [Pa] | 299 ± 14 | 150 ± 7 | 346 ± 16 | 1046 ± 77 |
| $E_2$ [Pa] | 289 ± 14 | 142 ± 7 | 320 ± 17 | 1024 ± 70 |
| $E_{inst}$ [Pa] | 1031 ± 47 | 479 ± 23 | 1175 ± 54 | 5029 ± 367 |
| $\tau_1$ [s] | 0.19 ± 0.01 | 0.19 ± 0.01 | 0.17 ± 0.01 | 0.19 ± 0.01 |
| $\tau_2$ [s] | 3.13 ± 0.14 | 2.93 ± 0.10 | 3.15 ± 0.15 | 3.23 ± 0.16 |
| $\eta_1$ [Pa s] | 56 ± 3 | 27 ± 1 | 58 ± 3 | 187 ± 14 |
| $\eta_2$ [Pa s] | 885 ± 55 | 403 ± 22 | 1052 ± 80 | 3198 ± 253 |

### Training self-organized maps with viscoelastic cell properties

In the next step self-organizing maps were trained with the scaled cell viscoelastic properties obtained after the different treatments. The performance of both iterative and batch algorithms was compared using a wide range of map parameters. These calculations were repeated multiple times to account for randomness in the initialisation and training. The maps shown from here on are the results of a batch trained map with 11 times 18 hexagonal nodes, a toroidal topography, a gaussian neighbourhood function, learning rates of (0.05, 0.01), Euclidean distance functions and 1000 iteration steps (map with smallest quantization error). Figure 1 shows the results of the SOM training.

**Figure 1.**
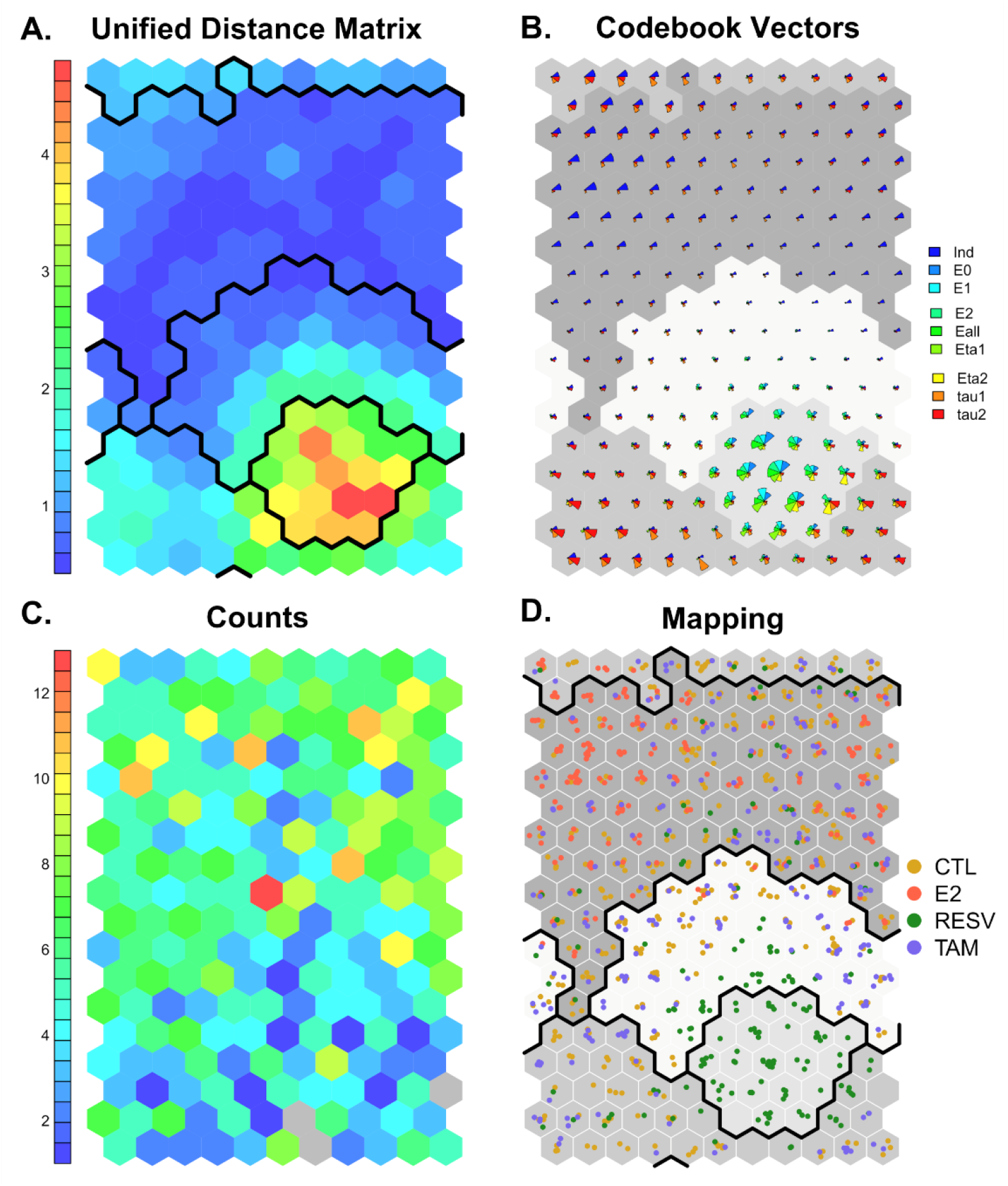
Results of training the self-organizing map using a batch algorithm. (A) Unified distance matrix plot. The colour scale corresponds to Euclidean distance between nodes. The post hoc clustering was performed using kmeans clustering with 4 centroids. (B) Fan-diagram showing the spatial distribution of the nine different variables on the 2D map. (C) Counts plot of the number of observations per node. In grey nodes, zero observations are placed. (D) Mapping of the treatment as input factors.

In Figure 1A the unified distance matrix of the SOM can be seen. The colour scale indicates the Euclidean distance between nodes, blue tones indicate small distances, while red tones indicate large ones. A distinct region with high dissimilarity can be observed at the bottom right of the map. Note that the map is toroidal. The dark black lines indicate the post hoc clustering of the nodes. Figure 1B shows the distributions of the single codebook vectors and the determined clusters in greyscales. This analysis can be used to analyse the distribution of the single input variables. Even for the low number of input variables, this analysis can be quite overwhelming. As an example, the nodes that showed high dissimilarity in Figure 1A on the right bottom side of the map showed large values of all moduli, while only possessing small indentations and intermediate relaxation times. Measurements with high indentation values appeared to be placed in vicinity on the top left-hand side of the map. Figure 1C shows the number of measurements that were placed in each node. Finally, Figure 1D uses the treatment factor (as these are known from the input data) and visualizes where measurements of respectively treated cells were placed on the map. The cluster on the bottom right side of the map is made up solely of resveratrol treated cells, while a large amount of estrogen treated cell measurements are located at the top left side of the map. Note that the number of clusters was defined as 4 to reflect the different treatments.

### Resveratrol and estrogen treated cells can be distinguished by unsupervised SOMs

Table 2 and 3 show a further analysis of the four clusters. Cluster 1 was predominantly (80%) made of cells from the control group and tamoxifen treated cells. Cluster 2 only included resveratrol treated cells. Cluster 3 was a mixture of all cell types, mostly control and tamoxifen treated ones. Finally, cluster 4 was made up mostly of control and estrogen treated cells. With respect to mechanics, cells placed in cluster 1 and 3 showed intermediate stiffnesses and indentations. Cluster 3 represented higher relaxation times than all other clusters. Resveratrol treated cells, located in Cluster 2, presented high stiffness, low indentations, high viscosities, and intermediate relaxation times. The characteristics of cluster 4 were very soft cells, with large indentations, low moduli, intermediate relaxation times and low viscosities. The cluster analysis shows that resveratrol treated cells can be readily distinguished from the other groups, while estrogen treated cells could mostly be found in a cluster together with control and tamoxifen treated cells. As expected from the mechanical properties, control cells and tamoxifen treated cells co-localised on the maps and the SOMs performed in this way were not able to distinguish between them.

**Table 2.** Analysis of the four cluster with respect to kind of treatment.

| Cluster | N | N (CTL) | N (E2) | N (RESV) | N (TAM) |
| --- | --- | --- | --- | --- | --- |
| 1 | 227 | 104 | 7 | 39 | 77 |
| 2 | 69 | 0 | 0 | 69 | 0 |
| 3 | 195 | 98 | 13 | 32 | 52 |
| 4 | 545 | 210 | 204 | 31 | 100 |

**Table 3.** Analysis of the four cluster with respect to the determined viscoelastic properties. The values shown are mean values ± the standard error.

|  | Cluster 1 | Cluster 2 | Cluster 3 | Cluster 4 |
| --- | --- | --- | --- | --- |
| $\delta$ [$\mu\text{m}$] | $1.60 \pm 0.02$ | $0.84 \pm 0.02$ | $2.04 \pm 0.04$ | $2.52 \pm 0.05$ |
| $E_{\infty}$ [Pa] | $987 \pm 69$ | $5348 \pm 207$ | $537 \pm 45$ | $275 \pm 13$ |
| $E_1$ [Pa] | $510 \pm 19$ | $1850 \pm 68$ | $352 \pm 15$ | $188 \pm 7$ |
| $E_2$ [Pa] | $498 \pm 19$ | $1773 \pm 55$ | $351 \pm 20$ | $175 \pm 8$ |
| $E_{inst}$ [Pa] | $1996 \pm 95$ | $8971 \pm 294$ | $1240 \pm 74$ | $638 \pm 26$ |
| $\tau_1$ [s] | $0.13 \pm 0.01$ | $0.18 \pm 0.01$ | $0.27 \pm 0.01$ | $0.17 \pm 0.01$ |
| $\tau_2$ [s] | $2.21 \pm 0.08$ | $3.13 \pm 0.13$ | $5.27 \pm 0.14$ | $2.71 \pm 0.09$ |
| $\eta_1$ [Pa s] | $71 \pm 4$ | $322 \pm 14$ | $94 \pm 4$ | $32 \pm 2$ |
| $\eta_2$ [Pa s] | $1124 \pm 60$ | $5291 \pm 232$ | $1859 \pm 133$ | $477 \pm 27$ |

**Table 3.** Analysis of the four cluster with respect to the determined viscoelastic properties. The values shown are mean values ± the standard error.
|  | Cluster 1 | Cluster 2 | Cluster 3 | Cluster 4 |
| --- | --- | --- | --- | --- |
| $\delta$ [μm] | 1.60 ± 0.02 | 0.84 ± 0.02 | 2.04 ± 0.04 | 2.52 ± 0.05 |
| $E_{\infty}$ [Pa] | 987 ± 69 | 5348 ± 207 | 537 ± 45 | 275 ± 13 |
| $E_1$ [Pa] | 510 ± 19 | 1850 ± 68 | 352 ± 15 | 188 ± 7 |
| $E_2$ [Pa] | 498 ± 19 | 1773 ± 55 | 351 ± 20 | 175 ± 8 |
| $E_{inst}$ [Pa] | 1996 ± 95 | 8971 ± 294 | 1240 ± 74 | 638 ± 26 |
| $\tau_1$ [s] | 0.13 ± 0.01 | 0.18 ± 0.01 | 0.27 ± 0.01 | 0.17 ± 0.01 |
| $\tau_2$ [s] | 2.21 ± 0.08 | 3.13 ± 0.13 | 5.27 ± 0.14 | 2.71 ± 0.09 |
| $\eta_1$ [Pa s] | 71 ± 4 | 322 ± 14 | 94 ± 4 | 32 ± 2 |
| $\eta_2$ [Pa s] | 1124 ± 60 | 5291 ± 232 | 1859 ± 133 | 477 ± 27 |

### SOMs for exploration of mechanical property interconnectivity

One strength of self-organizing maps is that they allow exploration of interconnectivity of different input variables. Figure 2 shows the component maps of the 9 input variables after training. In addition, the cluster analysis was superimposed on the maps. Note that some variables were plotted in logarithmic scale to allow for better comparison.

**Figure 2.**
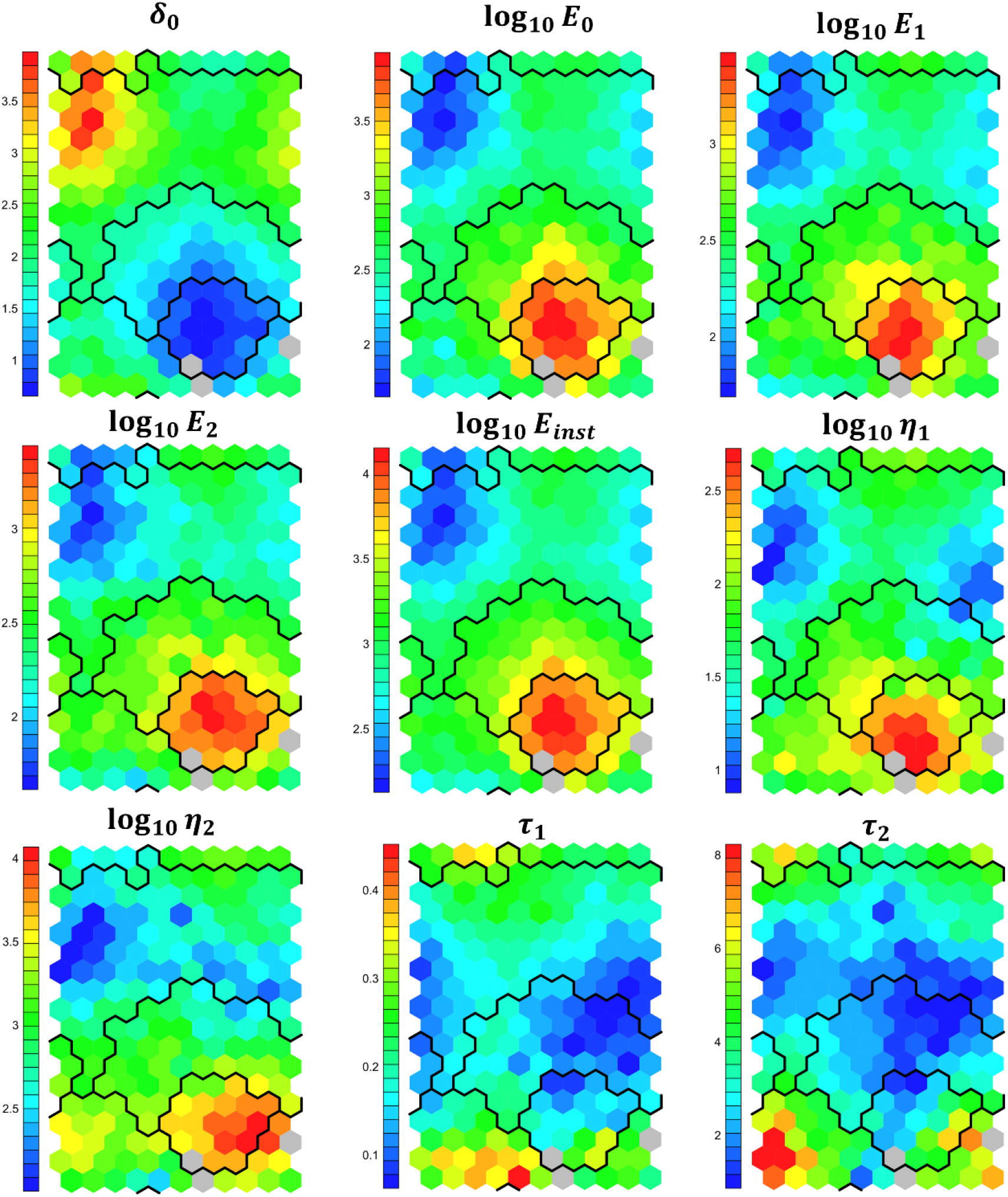
Component maps of the nine input parameters after training the SOMs. The colour coding indicates the magnitude of the nodes. Note that indentation and relaxation times are normal scaled with μm and s as units, while the moduli and viscosities are logarithmically scaled to allow for comparison. The black lines show the borders of the four calculated clusters. Grey nodes have no measurements placed in them.

A correlation between all moduli and the viscosities was observed. In addition, these six parameters were negatively correlated with the indentation. This fits together with the mechanical model, as cells that are softer should show lower moduli, higher fluidity (lower viscosity) and larger deformations. Two distinct regions of deformation (or moduli and viscosities) can be seen in the component maps: On the left top, a region with high indentation and low stiffness as well as viscosity was found, which corresponded mostly to estrogen treated cells. On the other hand, an island of low indentation, large moduli and large viscosities was detected on the bottom right part of the maps, which mostly displayed measurements made on resveratrol treated cells. Regarding the relaxation times, a large patch of small relaxation times was identified in the vertical middle region of the maps. Interestingly, these values appeared to correspond to intermediate deformation, moduli and viscosity values. A small patch of long relaxation times was noticed at the bottom left of the maps, which represented intermediate stiffness and deformation values. This indicates that the relaxation times are independent of the indentation and stiffnesses, and that viscosities are mostly determined by the moduli.

## Discussion

In this work we apply self-organizing maps to viscoelastic data of breast cancer cells derived from stress relaxation atomic force microscopy measurements after treatment with different ER interacting drugs. We employ an unsupervised approach and use the output layer of this artificial neural network for cluster analysis. We show that resveratrol treatment leads to stiffening and higher viscosities, estrogen treatment to a softening and fluidization, while tamoxifen treatment does not appear to significantly affect the mechanical properties. We then provide evidence that this type of analysis can potentially be used to classify cells according to treatment in an unsupervised manner.

Cancer cells show altered mechanical phenotypes compared to non-malignant counterparts ^16^. This results among other factors from changes in the mechanics of the environment, tumour hypoxia and aberrant gene expression, altering signalling pathways that affect cell metabolism, motility, and adhesion. Targeting the mechanical adaptation programme of cancer as part of therapy has been proposed, although its wide implications in cell function may lead to undesired side effects. Potential targets include disruption of actin filament assembly and dynamics, inhibition of myosin or targeting Rho/ROCK pathways, among others. Different drugs used as therapy such as paclitaxel, cisplatin, doxorubicin, and 5-fluorouracil, have been investigated for their effects on mechanical properties of cells from various origins. These treatments mostly led to cell stiffening, which contributed to altered cytoskeletal dynamics resembling a reversal of the epithelial to mesenchymal transition ^43–45^. Such changes are believed to lead to reduced cell migration, partly inhibiting cancer growth. We provide further evidence that resveratrol leads to stiffening of cells, with novel data regarding the viscoelastic properties, while estrogen leads to a softening and fluidization of the cells. Interestingly, tamoxifen does not appear to significantly alter the mechanics of MCF-7 cells on solid substrates. A more thorough analysis regarding cytoskeletal arrangements and protein expression patterns resulting from such treatments is needed.

In the present approach we have used viscoelastic data derived from stress relaxation measurements using a five-element Maxwell model fitting. We have arbitrarily omitted the use of any further characteristics of force spectroscopy curves on cells (such as hysteresis, work of adhesion, tethers, tension). In addition, we have decided to ignore the measurement data itself, but rather evaluate derived mechanical properties. A critical parameter to consider is the computational time the analysis takes. While the computation of the self-organizing map approach chosen is relatively straightforward, the data processing steps to calculate the viscoelastic properties from raw force-distance-curves is the time intensive step of this analysis (the used framework takes around 5 s for curve processing, fitting, and property calculation pre curve). Using simpler models (such as only elastic properties, indentations at a given force, hysteresis, adhesion properties) one can probably reduce the computational costs. In addition, we show that most of the variables used as inputs are correlated. Therefore, a priori reduction of data dimensionality using principal component analysis followed by statistically relevant principal components as input for the SOMs, will also speed up the analysis.

Most of the neural network or machine learning approaches that have been applied to AFM force spectroscopy data on biological materials are based on supervised methods. Recently, three different approaches have been provided in the literature: Using machine learning to classify the quality of curves ^25^, training a machine learning algorithm with curve shapes ^28^, or using curve characteristic properties as inputs for supervised training ^27,29,46^. Such approaches have shown promising results in correctly classifying cancer cells from healthy cells, as well as cancerous tissue. Compared to these approaches, SOMs arguably show strengths in data exploration, enabling the simplification of a multidimensional data space to 2D representations. In addition, SOMs can also be performed supervised. With respect to the ability of this SOM approach to identify the treatment of breast cancer cells, we report mixed results. The addition of further independent variables, including more measurements, and morphological characteristics of the cells will probably help in training a SOM algorithm to successfully classify the different types of cells unsupervised.

## Materials and Methods

Figure 3 shows the overall methodology performed in this study. Briefly, in step 1, cell mechanical properties were quantified using stress relaxation measurements via AFM. A scheme of such analysis can be seen in Figure 3B. A 5-element Maxwell model was fitted to the stress relaxation curves, resulting in 9 fitting parameters (Figure 3C). These mechanical parameters were used to train SOMs (Figure 3D). Finally, these SOMs were used to visualize the result of the artificial neural network. To test the ability of the trained aNNs to distinguish between cells with different mechanical properties, MCF-7 breast cancer cells were treated with different substances used in breast cancer therapy that are known to influence cell mechanical properties.

**Figure 3.**
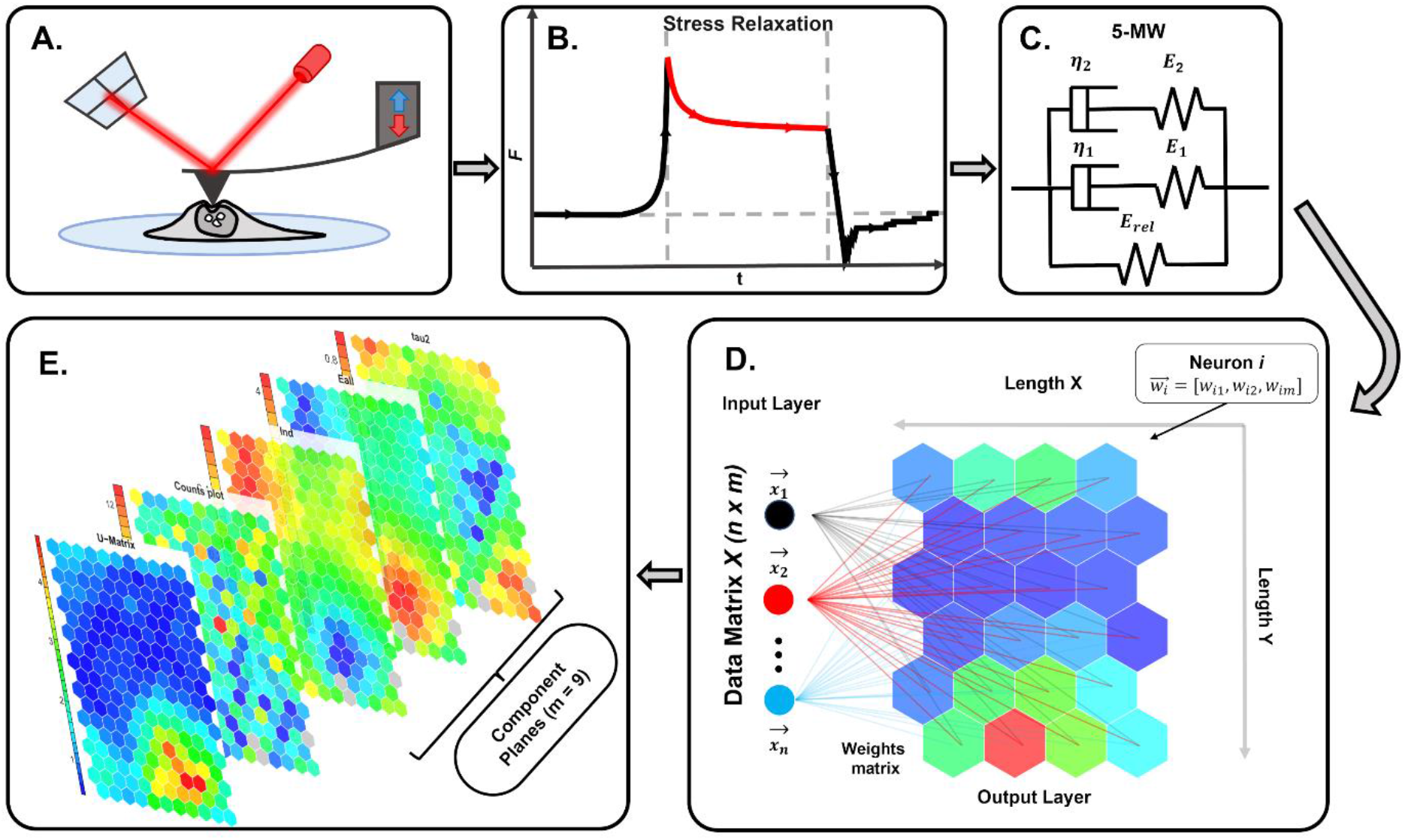
Methodology of the present study. A. Cell viscoelastic properties are measured using stress relaxation experiments with AFM. B and C. The resulting stress relaxation segment (force decay over time while keeping the deformation constant) is fitted using a five-element Maxwell model. This results in 9 input variables. D. A self-organizing map is trained unsupervised. E. The complex interconnectivity of the data can be represented in 2D maps, allowing for data exploration.

### Cell mechanical measurements

MCF-7 cells were obtained from the American Type Culture Collection (ATCC) and grown as previously described ^47^. Cells were seeded at a concentration of 200,000 cells/mL on plasma-treated glass slides in DMEM medium supplemented with 10% fetal bovine serum (FBS) and 1% penicillin/streptomycin at 37 °C in 5% CO_2_. Cells were either treated for 48 hours with DMSO (0.05 %), 100 nM estrogen (E2), 50 μM resveratrol (RESV) or 10 μM tamoxifen (TAM). For mechanical measurements, a JPK Nanowizard III with a CellHesion extension was used. Pyramidal cantilevers (DNP-S, B, Bruker), with nominal stiffness of 0.12 N/m, a resonance frequency of 23 kHz in air, an opening angle of 22 ° and a nominal tip radius of 10 nm were used. Calibration by thermal noise making use of the equipartition theorem was performed for each cantilever ^48^. Measurements were done with an approach and retract rate of 5 μm/s, a maximum load of 1 nN, curve lengths of 50 μm, a constant deformation 10 s pause segment at an initial load of 1 nN and a sampling rate of 1024 Hz. Measurements were done in L15 medium at 37 °C.

### Data evaluation

Data was extracted using the JPKSPM software (JPK, Bruker), and all further steps were performed in R. Data pre-processing was done making use of the R afmToolkit, a package for AFM force-distance and force-time analysis developed by our group ^49–51^. Briefly, contact and detachment points were calculated, the curves were corrected for their baselines and the sample deformation (*δ*) was determined. In the present analysis, only the stress relaxation segments were considered. Those were fitted with a five-element Maxwell model as

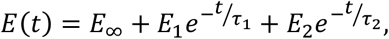

Where *E*(*t*) is the relaxation modulus, *E*_∞_ the equilibrium modulus, *E*_1_ and *E*_2_ the moduli of the springs in Maxwell arms and *τ*_1_ and *τ*_2_ the relaxation times of the dashpots. The viscosity *η*_*i*_ of the dashpots is defined as

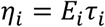

### Self-organizing maps

Self-organizing maps were first introduced by Kohonen and have since received more and more attention in wider fields of statistics, chemometrics and life sciences. Their major use is found in data exploration and visualization, as they enable the breakdown of complex data matrices and the interaction of components via 2D similarity representations. In addition, the representation keeps the topological information of the input data. Here we describe the principles of the classical iterative and the batch algorithms. Any reader is referred to the extensive reviews on the theoretical backgrounds, application and usage of SOMs in statistical analysis, life sciences and industry ^35,36,52^. Such applications are wide ranged, including analysis of spectroscopy data ^53,54^, automated feature extraction from microscopy images ^55^, segmentation and grading of tumours or application to omics data ^56,57^.

SOMs are artificial neural networks that use unsupervised training procedures (competition, cooperation, and adaptation). In principle, a (mostly) 2D network of neurons (nodes, codebooks) is set up that is trained to represent the distance structure of the input data as closely as possible (see Figure XYZ D). In the case of SOMs, nodes are defined by codebook vectors (weights) that encompass all variables of the input data. Prior to training, the network is initialized either randomly or linearly. One needs to a priori define network properties such as topology (hexagonal or rectangular), architecture of the grid (dimensions, shape, repetitiveness, dynamically growing SOMs), number of neurons (depends on input data, approach, and analysis route). Depending on the type of distance measure applied, input data needs to be normalized to account for different scaling.

In the original iterative approach, the following training steps are performed:

1. Randomly select an input vector.
2. Calculate the distance between the input and all neurons in the network (using Euclidean distance after normalization works for most cases).
3. Determine the best matching unit (BMU) with the lowest distance to the input vector (competition).
4. Calculate the neighbouring neurons that will also be updated according to the chosen neighbourhood function (gaussian, bubble) (cooperation)
5. Update the BMU and the neighbours applying learning rates, by moving them closer to the input vector (adaptation).
6. Repeat steps 1 to 6 until a given number of iterations is reached or the system converges to stability.

Both the neighbourhood distance and the learning rate are decreasing monotonically with each iteration. The learning rate is adjusted depending on the proximity of the neuron to the BMU. The learning phase is split in two main segments, first with a high learning rate (ordering phase) and then with a lower one (convergence phase). In batch SOMs, a similar approach is performed but rather all neurons are compared with all input data.

### Application of SOMs to breast cancer viscoelastic properties

As input data, we used the 9 properties (*δ, E*_∞_, *E*_1_, *E*_2_, *E*_*inst*_, *τ*_1_, *τ*_2_, *η*_1_, *and η*_2_) derived from the fitting of the stress relaxation segments. Outliers were removed and data was normalized. For SOM calculations, the R package Kohonen was used. Rectangular, toroidal maps with hexagonal nodes were used. The x-y-ratio of the maps was determined from the relationship of the eigenvalues of the first two principal components of the data. A Gaussian neighbourhood function and Euclidean distance functions were utilized. The learning rate was set to 0.05 in the beginning and decreased to 0.01. The neighbourhood function was defined as exponential decay starting with a value that covered 2/3 of all unit-to-unit distances. For the classical, iterative approaches, 50,000 iterations were performed, the calculations were repeated 50 times and the map with the lowest quantization error was chosen. Similarly, batch SOMs were performed with 1,000 iterations and repeated 50 times. The results shown in this manuscript correspond to the batch SOM with the lowest quantization error.

## Acknowledgements

The authors want to acknowledge Amsatou Andorfer-Sarr for maintenance of cell culture facilities and Barbara Zbiral for critical discussion of the data and manuscript. This research was partially funded by Austrian Science Fund (FWF), grant number P35777-B and by the Spanish Ministry of Science and Innovation (PID2020-118464RB-I00) and Elkartek funding (KK-2020/00008) by the Basque Government (MdMV).

## Author contributions (names must be given as initials)

AW: Conceptualization, Methodology, Software, Formal Analysis, Investigation, Writing – Original Draft, Writing – review & editing, visualization. MdM: Writing – review & editing, supervision, project administration. JLTH: Conceptualization, Writing – Original Draft, Writing – review & editing, visualization, supervision, project administration, funding acquisition.

## Data availability statement (mandatory)

Data is available per request from the corresponding author.

## Additional Information (including a Competing Interests Statement)

The authors declare no competing interests.

## References

1. Vogel, V. & Sheetz, M. Local force and geometry sensing regulate cell functions. Nat. Rev. Mol. Cell Biol. 7, 265–75 (2006).

2. Vining, K. H. & Mooney, D. J. Mechanical forces direct stem cell behaviour in development and regeneration. Nat. Rev. Mol. Cell Biol. 18, 728–742 (2017).

3. Roca-Cusachs, P., Conte, V. & Trepat, X. Quantifying forces in cell biology. Nat. Cell Biol. 19, 742–751 (2017).

4. Petridou, N. I., Spiró, Z. & Heisenberg, C.-P. Multiscale force sensing in development. Nat. Cell Biol. 19, 581–588 (2017).

5. Huse, M. Mechanical forces in the immune system. Nat. Rev. Immunol. 17, 679–690 (2017).

6. Romani, P., Valcarcel-Jimenez, L., Frezza, C. & Dupont, S. Crosstalk between mechanotransduction and metabolism. Nat. Rev. Mol. Cell Biol. 22, 22–38 (2021).

7. Winkler, J., Abisoye-Ogunniyan, A., Metcalf, K. J. & Werb, Z. Concepts of extracellular matrix remodelling in tumour progression and metastasis. Nat. Commun. 11, 5120 (2020).

8. Wirtz, D., Konstantopoulos, K. & Searson, P. C. The physics of cancer: the role of physical interactions and mechanical forces in metastasis. Nat. Rev. Cancer 11, 512–522 (2011).

9. Fife, C. M., McCarroll, J. A. & Kavallaris, M. Movers and shakers: Cell cytoskeleton in cancer metastasis. Br. J. Pharmacol. 171, 5507–5523 (2014).

10. Lekka, M. et al. Cancer cell recognition – Mechanical phenotype. Micron 43, 1259–1266 (2012).

11. Lin, H.-H. et al. Mechanical phenotype of cancer cells: cell softening and loss of stiffness sensing. Oncotarget 6, 20946–20958 (2015).

12. Brill-Karniely, Y. et al. Triangular correlation (TrC) between cancer aggressiveness, cell uptake capability, and cell deformability. Sci. Adv. 6, eaax2861 (2022).

13. Di Carlo, D. A mechanical biomarker of cell state in medicine. J. Lab. Autom. 17, 32–42 (2012).

14. Northcott, J. M., Dean, I. S., Mouw, J. K. & Weaver, V. M. Feeling Stress: The Mechanics of Cancer Progression and Aggression. Frontiers in Cell and Developmental Biology vol. 6 17 (2018).

15. Kozminsky, M. & Sohn, L. L. The promise of single-cell mechanophenotyping for clinical applications. Biomicrofluidics 14, 31301 (2020).

16. Nguyen, L. T. S., Jacob, M. A. C., Parajón, E. & Robinson, D. N. Cancer as a biophysical disease: Targeting the mechanical-adaptability program. Biophys. J. 121, 3573–3585 (2022).

17. Moeendarbary, E. & Harris, A. R. Cell mechanics: Principles, practices, and prospects. Wiley Interdiscip. Rev. Syst. Biol. Med. 6, 371–388 (2014).

18. Wu, P.-H. et al. A comparison of methods to assess cell mechanical properties. Nat. Methods 15, 491–498 (2018).

19. Efremov, Y. M., Okajima, T. & Raman, A. Measuring viscoelasticity of soft biological samples using atomic force microscopy. Soft Matter 16, 64–81 (2019).

20. Lekka, M. et al. Elasticity of normal and cancerous human bladder cells studied by scanning force microscopy. Eur. Biophys. J. 28, 312–316 (1999).

21. Plodinec, M. et al. The nanomechanical signature of breast cancer. Nat. Nanotechnol. 7, 757–765 (2012).

22. Azuri, I., Rosenhek-goldian, I., Regev-rudzki, N., Fantner, G. & Cohen, S. R. The role of convolutional neural networks in scanning probe microscopy : a review. Beilstein J. Nanotechnol. 878–901 (2021) doi:10.3762/bjnano.12.66.

23. Sotres, J., Boyd, H. & Gonzalez-Martinez, J. F. Enabling autonomous scanning probe microscopy imaging of single molecules with deep learning. Nanoscale 13, 9193–9203 (2021).

24. Braunsmann, C. & Schäffer, T. E. Note: Artificial neural networks for the automated analysis of force map data in atomic force microscopy. Rev. Sci. Instrum. 85, 56104 (2014).

25. Müller, P. et al. nanite: using machine learning to assess the quality of atomic force microscopy-enabled nano-indentation data. BMC Bioinformatics 20, 465 (2019).

26. Ilieva, N. I., Galvanetto, N., Allegra, M., Brucale, M. & Laio, A. Automatic classification of single-molecule force spectroscopy traces from heterogeneous samples. Bioinformatics 36, 5014–5020 (2020).

27. Sotres, J., Boyd, H. & Gonzalez-Martinez, J. F. Locating critical events in AFM force measurements by means of one-dimensional convolutional neural networks. Sci. Rep. 12, 12995 (2022).

28. Minelli, E. et al. A fully-automated neural network analysis of AFM force-distance curves for cancer tissue diagnosis. Appl. Phys. Lett. 111, 143701 (2017).

29. Sokolov, I. et al. Noninvasive diagnostic imaging using machine-learning analysis of nanoresolution images of cell surfaces: Detection of bladder cancer. Proc. Natl. Acad. Sci. 115, 12920–12925 (2018).

30. Ciasca, G. et al. Efficient Spatial Sampling for AFM-Based Cancer Diagnostics: A Comparison between Neural Networks and Conventional Data Analysis. Condensed Matter vol. 4 (2019).

31. Prasad, S. et al. Atomic Force Microscopy Detects the Difference in Cancer Cells of Different Neoplastic Aggressiveness via Machine Learning. Adv. NanoBiomed Res. 1, 2000116 (2021).

32. Tian, Y., Lin, W., Qu, K., Wang, Z. & Zhu, X. Insights into cell classification based on combination of multiple cellular mechanical phenotypes by using machine learning algorithm. J. Mech. Behav. Biomed. Mater. 128, 105097 (2022).

33. Kohonen, T. Self-organized formation of topologically correct feature maps. Biol. Cybern. 43, 59–69 (1982).

34. Kohonen, T., Oja, E., Simula, O., Visa, A. & Kangas, J. Engineering applications of the self-organizing map. Proc. IEEE 84, 1358–1384 (1996).

35. Brereton, R. G. Self organising maps for visualising and modelling. Chem. Cent. J. 6 Suppl 2, S1 (2012).

36. Kohonen, T. Essentials of the self-organizing map. Neural Networks 37, 52–65 (2013).

37. Sung, H. et al. Global Cancer Statistics 2020: GLOBOCAN Estimates of Incidence and Mortality Worldwide for 36 Cancers in 185 Countries. CA. Cancer J. Clin. 71, 209–249 (2021).

38. Bouris, P. et al. Estrogen receptor alpha mediates epithelial to mesenchymal transition, expression of specific matrix effectors and functional properties of breast cancer cells. Matrix Biol. 43, 42–60 (2015).

39. Padilla-Rodriguez, M. et al. The actin cytoskeletal architecture of estrogen receptor positive breast cancer cells suppresses invasion. Nat. Commun. 9, 2980 (2018).

40. Iturri, J. et al. Resveratrol-induced temporal variation in the mechanical properties of MCF-7 breast cancer cells investigated by atomic force microscopy. Int. J. Mol. Sci. 20, (2019).

41. Bischoff, P. et al. Estrogens Determine Adherens Junction Organization and E-Cadherin Clustering in Breast Cancer Cells via Amphiregulin. iScience 23, 101683 (2020).

42. Zbiral, B., Weber, A., Iturri, J., Vivanco, M. d. M. & Toca-Herrera, J. L. Estrogen Modulates Epithelial Breast Cancer Cell Mechanics and Cell-to-Cell Contacts. Materials vol. 14 (2021).

43. Raudenska, M. et al. Cisplatin enhances cell stiffness and decreases invasiveness rate in prostate cancer cells by actin accumulation. Sci. Rep. 9, 1660 (2019).

44. Rodríguez-Nieto, M. et al. Viscoelastic properties of doxorubicin-treated HT-29 cancer cells by atomic force microscopy: the fractional Zener model as an optimal viscoelastic model for cells. Biomech. Model. Mechanobiol. 19, 801–813 (2020).

45. Kubiak, A., Zieliński, T., Pabijan, J. & Lekka, M. Nanomechanics in Monitoring the Effectiveness of Drugs Targeting the Cancer Cell Cytoskeleton. Int. J. Mol. Sci. 21, (2020).

46. Darling, E. M., Topel, M., Zauscher, S., Vail, T. P. & Guilak, F. Viscoelastic properties of human mesenchymally-derived stem cells and primary osteoblasts, chondrocytes, and adipocytes. J. Biomech. 41, 454–464 (2008).

47. Iriondo, O. et al. Distinct breast cancer stem/progenitor cell populations require either HIF1α or loss of PHD3 to expand under hypoxic conditions. Oncotarget 6, 31721–31739 (2015).

48. Butt, H.-J. & Jaschke, M. Calculation of thermal noise in atomic force microscopy. Nanotechnology 6, 1–7 (1995).

49. Benítez, R., Moreno-flores, S., Bolós, V. J. & Toca-Herrera, J. L. A new automatic contact point detection algorithm for AFM force curves. Microsc. Res. Tech. 76, 870–876 (2013).

50. Benítez, R., Bolós, V. J. & Toca-Herrera, J. L. afmToolkit: an R Package for Automated AFM Force-Distance Curves Analysis. R J. 9, 291–308 (2017).

51. Weber, A., Benitez, R. & Toca-Herrera, J. L. Measuring (biological) materials mechanics with atomic force microscopy. 4. Determination of viscoelastic cell properties from stress relaxation experiments. Microsc. Res. Tech. n/a, (2022).

52. Lloyd, G. R., Brereton, R. G. & Duncan, J. C. Self Organising Maps for distinguishing polymer groups using thermal response curves obtained by dynamic mechanical analysis. 133, 1046–1059 (2008).

53. Beckonert, O., Monnerjahn, J., Bonk, U. & Leibfritz, D. Visualizing metabolic changes in breast-cancer tissue using 1H-NMR spectroscopy and self-organizing maps. NMR Biomed. 16, 1–11 (2003).

54. Riese, F. M., Keller, S. & Hinz, S. Supervised and Semi-Supervised Self-Organizing Maps for Regression and Classification Focusing on Hyperspectral Data. Remote Sensing vol. 12 (2020).

55. Kang, M.-S., Kim, H.-R. & Kim, M.-H. Cell Classification in 3D Phase-Contrast Microscopy Images via Self-Organizing Maps BT - Advances in Visual Computing. in (eds. Bebis, G.et al.) 652–661 (Springer International Publishing, 2014).

56. Vijayakumar, C., Damayanti, G., Pant, R. & Sreedhar, C. M. Segmentation and grading of brain tumors on apparent diffusion coefficient images using self-organizing maps. Comput. Med. imaging Graph. Off. J. Comput. Med. Imaging Soc. 31, 473–484 (2007).

57. Binder, H. et al. Integrated Multi-Omics Maps of Lower-Grade Gliomas. Cancers (Basel). 14, (2022).

